# What happens with bacteria in the roots affects what happens with fungi in the leaves

**DOI:** 10.1101/2023.12.08.570838

**Authors:** Rachelle Fernández-Vargas, Fabiana Tabash-Porras, Keilor Rojas-Jiménez

## Abstract

The impact of plant-bacteria symbioses in roots on plant-fungus interactions in distant tissues, such as leaves, remains a significant knowledge gap, particularly for tropical ecosystems. We employed amplicon metabarcoding of ITS2 sequences to analyze the endophytic fungal communities of eight tropical Fabaceae tree species to determine variations according to the host’s root nodulation capacity and the host’s subfamily affiliation. Nitrogen-fixing bacteria in root nodules significantly impacted the diversity and composition of fungal endophytes, while subfamily affiliation had a less pronounced effect. In non-nodulating plants, leaves harbored a much larger diversity of fungi, with 25 fungal classes and 72 orders compared to 17 classes and 45 orders in nodulating plants. Notably, 40% of the orders were unique to non-nodulating plants, whereas only 5% were exclusive to nodulating species. This pattern was further reflected in the number of fungal ASVs, with non-nodulating plants averaging 130.5 compared to 42.7 in nodulating plants. Similarly, Shannon diversity was significantly higher in non-nodulating plants. This study demonstrates that interactions between Fabaceae plants and nitrogen-fixing root bacteria significantly influence the richness and diversity of the fungal community present in leaves. Our findings contribute valuable insights into the complex interplay between plants and microorganisms, particularly in the context of tropical Fabaceae tree species and their endophytic fungi.

## 1. Introduction

The legume family (Fabaceae) is one of the most diverse plant groups in the world, playing a crucial role in both agriculture and forest ecological dynamics (Azani et al., 2017; Lewis, 2005; Põlme et al., 2020). The remarkable ecological success of legumes, particularly in tropical ecosystems, can be attributed to their ability to form symbiotic relationships with nitrogen-fixing bacteria (de Bedout-Mora et al., 2022; Mathesius, 2022; Parker, 2008). This symbiotic relationship provides legumes with a vital source of nitrogen, enhancing their adaptation capacity and contributing to their abundance and species diversity (James, 2022; Mathesius, 2022; Sprent et al., 2017). The symbiotic interaction between the host organism and the symbiont manifests itself through the formation of specialized root structures called nodules, where nitrogen fixation occurs (de Faria et al., 2022; Ferguson et al., 2019).

By acquiring nitrogen through symbiotic bacteria, nitrogen-fixing leguminous plants gain a significant competitive advantage over non-fixing species. These symbiotic bacteria convert atmospheric nitrogen into plant-soluble compounds, such as ammonia, which the host plant can readily utilize. Ammonia then enters the food chain as it becomes incorporated into amino acids and proteins (Adams et al., 2016; Ferguson et al., 2019, 2010; Sprent, 2008). This process not only influences nutrient availability for plants but also affects their interactions with other endosymbionts, including endophytic fungi (Danko et al., 2021; Franklin et al., 2020; Põlme et al., 2018a).

Endophytic fungi are microorganisms that live in association with plants for most or all of their life cycle, primarily residing within the leaves, stems, and roots of their hosts (Clay and Schardl, 2002; Schulz and Boyle, 2005). These fungi are considered crucial symbiotic partners for plant development, forming a mutually beneficial relationship where they receive nutrients and protection from their host in exchange for enhancing plant growth, increasing stress tolerance, and producing a diverse array of secondary metabolites that protect against herbivores and phytopathogenic microorganisms (Alam et al., 2021; Arnold et al., 2003; Rodriguez et al., 2009).

The compatibility between a host plant and its endophytic fungus can be influenced by environmental factors, a phenomenon known as ecological specificity. Moreover, the colonization and diversity of endophytic fungi within different plant tissues can exhibit noticeable variation, which is primarily attributed to biotic factors (Põlme et al., 2018b; Rojas-Jimenez et al., 2016; Rojas-Jimenez and Tamayo-Castillo, 2021). The connection between plants and endophytic fungi is not a random occurrence but rather represents a complex symbiotic relationship that has evolved (dos Reis et al., 2022; Reis, 2022). The colonization of plant tissues by endophytic fungi is deeply influenced by the specificity and specialization inherent to this symbiotic relationship. These factors, in turn, depend on a series of intrinsic characteristics present in both plants and fungi to ensure the successful establishment of symbiosis (dos Reis et al., 2022; Põlme et al., 2018b).

The disparities in host preference among endophytic fungi arise from multiple determinants, such as the closeness of the association, phylogenetic differences, competitive interactions, and resource allocation between symbionts. Together, these factors form a complex network that shapes the dynamics of colonization and the diversity of endophytic fungi in plant tissues, particularly in leaves (Heilmann-Clausen et al., 2016; Molina, 1992).

We hypothesize that the composition of endophytic fungal communities of leguminous tropical trees will differ depending on whether or not the host plant can establish symbiotic relationships with nitrogen-fixing bacteria. Specifically, we predict that leguminous plants capable of forming root nodules with nitrogen-fixing bacteria will exhibit lower diversity in their leaf mycobiome compared to those that do not engage in this type of symbiosis.

This prediction stems from the observation that microbial ecosystems with higher levels of disturbance or nutrient richness tend to be dominated by a few key species, whereas more constrained environments support a greater diversity of microorganisms, which likely cooperate to handle these challenging conditions. In this regard, our study delves into the intricate relationships between plant tissues and microbial communities in tropical leguminous trees, providing valuable insights into their dynamic interactions. Specifically, it investigates the interplay between nitrogen-fixing bacteria residing in plant roots and their influence on the fungal communities inhabiting the leaves. These findings hold significant practical implications for both ecosystem management and sustainable forestry practices.

## 2. Materials and Methods

### Sample collection

We conducted our research with eight plant species from the Fabaceae family, each with three biological replicates. Four of these species produce root nodules while four do not. Furthermore, four species belong to the Papilionoideae subfamily, and four to the Caesalpinioideae subfamily (**Table 1**). We used three replicates for each tree species. We obtained the specimens (50-70 cm tall) from a plant nursery located at the Santa Ana Conservation Center, San Jose, Costa Rica. After purchase, the plants were subjected to a three-month acclimatization period on the patio of a house in Barva de Heredia (temperature range 18-28°C, altitude 1100 m). The plants were irrigated daily. No pesticides were used during this period. This research project was conducted in full compliance with the ethical standards and regulations set forth by the Institutional Biodiversity Commission at the University of Costa Rica (permit VI-4056-2020).

**Table 1.** List of Fabaceae tree species selected for this study. The information on the taxonomy assignation to the subfamily level and the ability to form root nodules is included.

| Sample | Species | Subfamily | Nodulation |
| --- | --- | --- | --- |
| P1 | *Dalbergia retusa* | Papilionoideae | Positive |
| P2 | *Platymiscium pinnatum* | Papilionoideae | Positive |
| C1 | *Enterolobium cyclocarpum* | Caesalpinioideae | Positive |
| C2 | *Leucaena leucocephala* | Caesalpinioideae | Positive |
| P3 | *Dipteryx panamensis* | Papilionoideae | Negative |
| P4 | *Myrospermum frutescens* | Papilionoideae | Negative |
| C3 | *Senna reticulata* | Caesalpinioideae | Negative |
| C4 | *Cassia grandis* | Caesalpinioideae | Negative |

We collected approximately 250 mg of leaf tissues from each plant. We performed a superficial cleaning of the leaves using distilled water, we cut the healthy and undamaged leaves into approximately 5 mm pieces and placed them into Eppendorf tubes. We then conducted three consecutive washes using autoclaved and distilled water. The tissues were rinsed with 70% ethanol for 1 minute, followed by three additional rinses using autoclaved distilled water. Sterilization was achieved using a 2% sodium hypochlorite solution for 3 minutes, followed by five 1-minute rinses in distilled water. Following the sterilization process, we pulverized the plant material with liquid nitrogen to facilitate the extraction of genomic DNA.

### Molecular analyses

For DNA extraction, we employed the Qiagen PowerSoil® DNA extraction kit (Qiagen, Carlsbad, CA, USA) following the manufacturer’s protocols. We quantified DNA concentrations at 260 nm using a Nanodrop (Nanodrop 2000c spectrophotometer, Thermo Scientific) and stored the samples at -20°C. Subsequently, we constructed the eukaryotic amplicon library based on the ITS region by utilizing primers for amplifying fungal endophytes ITS1-F1-F (CTTGGTCATTTAGAGGAAGTAA) and ITS1-F1-R (GCTGCGTTCTTCATCGATGC). We conducted PCR amplification of target regions with specific primers linked to barcodes and PCR products with the appropriate length were selected via 1% agarose gel electrophoresis. The libraries were sequenced on an Illumina paired-end platform to generate 250-base pair paired-end reads (Illumina Novaseq, Novogene Bioinformatics Technology Co., Ltd, CA, USA).

### Bioinformatic Analysis

We conducted the bioinformatic analysis using DADA2 (version 1.26) to process the fastq format files obtained from Illumina paired-end sequencing. Additionally, we generated a table of Amplicon Sequence Variants (ASVs), which provide higher resolution equivalents compared to conventional OTUs (Callahan et al., 2016). We removed primers and adapters, inspected read quality, filtered and trimmed sequences with a quality score below 30, assessed error rates, modeled and corrected amplicon errors, and inferred sequence variants. Subsequently, we merged forward and reverse reads to obtain noise-free complete sequences, removed chimeras, and constructed the ASV table. Taxonomy was assigned to ASVs using a training set of reference sequences with known taxonomy from the UNITE database version 9.0 (https://doi.plutof.ut.ee/doi/10.15156/BIO/2938067). We verified the consistency of taxonomic assignments and manually curated taxonomic consistency using the NCBI Genbank BLAST tool. Sequences from host plants were excluded. Sequence data were deposited at the NCBI Sequence Read Archive under accession number **PRJNA1049775** (https://www.ncbi.nlm.nih.gov/sra/PRJNA1049775).

### Statistical analysis

We conducted the statistical analysis using the R (R Core Team, 2023) and its interface, RStudio. We employed the statistical library Vegan v2.6-4 (Oksanen et al., 2022) to calculate alpha diversity and conduct a non-parametric multidimensional scaling analysis (NMDS). Abundance tables of ASVs were adjusted to express them as relative abundances and then transformed into a Bray-Curtis dissimilarity matrix. To determine significant differences in fungal community diversity based on nodulation presence, we implemented a non-parametric multivariate analysis of variance (PERMANOVA). Differences in the relative proportion of fungi and diversity indices based on subfamily and nodulation capacity were estimated using the Wilcoxon signed-rank test.

## 3. Results

The fungal community of the foliar samples was composed of 1.335 amplicon sequence variants (ASVs), according to the analysis of the ITS sequences. The fungal sequences were assigned to 7 phyla, 26 classes, and 76 orders. The most prevalent phyla were Ascomycota, comprising 86.8% of the total number of sequences, and Basidiomycota representing 13.1% of the sequences. Other less prevalent fungal phyla included Mortierellomycota and Chytridiomycota. Ascomycota was represented by 10 classes and 42 orders, while Basidiomycota was composed of 11 classes and 29 orders.

Fungal endophyte communities exhibited variations within samples but were primarily influenced by the host’s root nodulation capacity (**Fig. 1**). Notably, leaves of plant species that do not form nodules on their roots harbored a larger number of fungal taxa compared to nodulating species. Specifically, 25 fungal classes and 72 orders were identified in non-nodulating plants, while only 17 classes and 45 orders were found in nodulating plants.

**Fig. 1.**
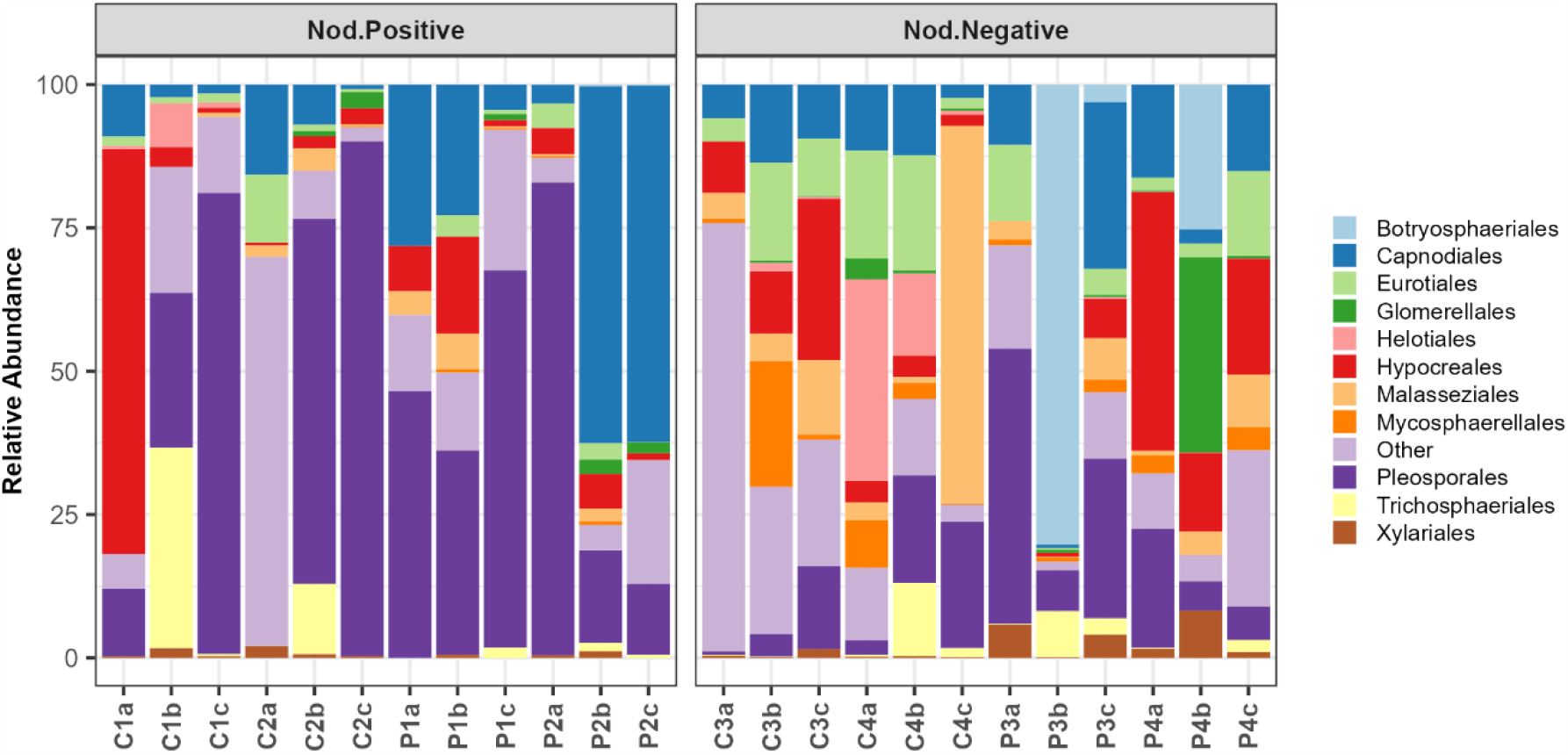
Relative abundance of the main fungal orders detected in leaves of eight tropical Fabaceae tree species. Panels separate species according to their capacity to form root nodules.

The fungal classes with the highest relative abundance included Dothideomycetes (56.3%), Sordariomycetes (16.9%), Eurotiomycetes (7.0%), and Malasseziomycetes (6.3%). Within Dothideomycetes the most abundant orders were Pleosporales (32.6%), Capnodiales (10.6%), and Botryosphaeriales (10.4%). Within Eurotiomycetes and Sordariomycetes the most abundant orders were Eurotiales (7.0%) and Hypocreales (6.9%), respectively.

The orders Capnodiales and Pleosporales were highly abundant in plants positive and negative for nodulation, however, we detected a larger proportion in leaves with positive nodulation. The most abundant genus within Capnodiales was *Cladosporium*, while *Preussia* and *Corynespora* were the most abundant genera within Pleosporales. The orders Botryosphaeriales, Eurotiales, Helotiales, Hypocreales, Malasseziales, Mycosphaerellales, and Xylariales were highly abundant in leaves of plants that do not form nodules. Botryosphaeriales and the genus *Phyllosticta* were detected only in plants with negative nodulation. The genera *Aspergillus, Engyodontium, Fusarium*, and *Malassezia* were among the genera with an elevated prevalence in leaves of negative nodulation.

The presence of nitrogen-fixing bacteria in root nodules significantly impacted the alpha diversity of fungal endophytes (**Fig. 2A**). However, this influence was less pronounced when considering subfamily affiliation (**Fig. 2B**). Plants lacking nodulation displayed significantly higher richness (Wilcoxon test, P < 0.05), with an average of 130.5 ASVs compared to 42.7 ASVs in nodulating plants. Similarly, Shannon diversity was significantly higher (Wilcoxon test, P < 0.05) in non-nodulating plants, with average values of 2.7 compared to 1.9 in nodulating plants. This trend persisted when analyzed by species (**Fig. 2C**).

**Fig. 2.**
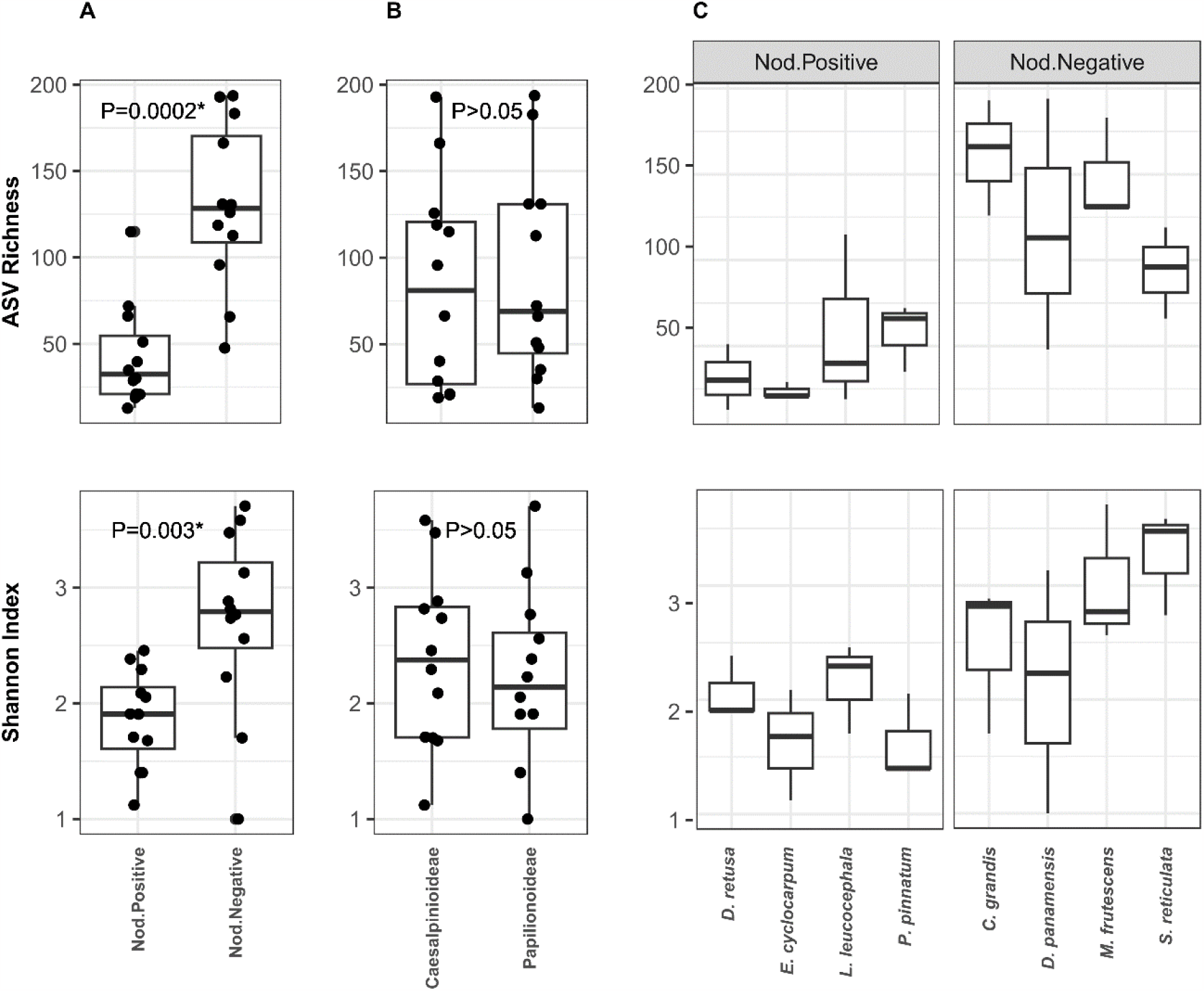
Alpha diversity values of foliar fungal communities according to nodulation capacity (Panel A), the Fabaceae subfamily (Panel B), and species (Panel C). The upper panels illustrate the richness of Amplicon Sequence Variants (ASVs), while the lower panels display the values of the Shannon Index. Statistically significant differences (Wilcox, P<0.05) between treatments are marked with an asterisk.

The NMDS analysis revealed distinct clustering patterns of endophyte communities based on the plant’s nodulation capacity (**Fig. 3A**). The PERMANOVA further confirmed significant differences in the structure of fungal communities between nodulating and non-nodulating plants (PERMANOVA, p = 0.004). Consistent with these results, the analysis of fungal orders in leaves showed that the majority (54.5%) were shared between the different species of leguminous trees studied. However, approximately 40% of the orders were found exclusively in non-nodulating plants, while only 5% were unique to nodulating species (**Fig. 3B**).

**Fig. 3.**
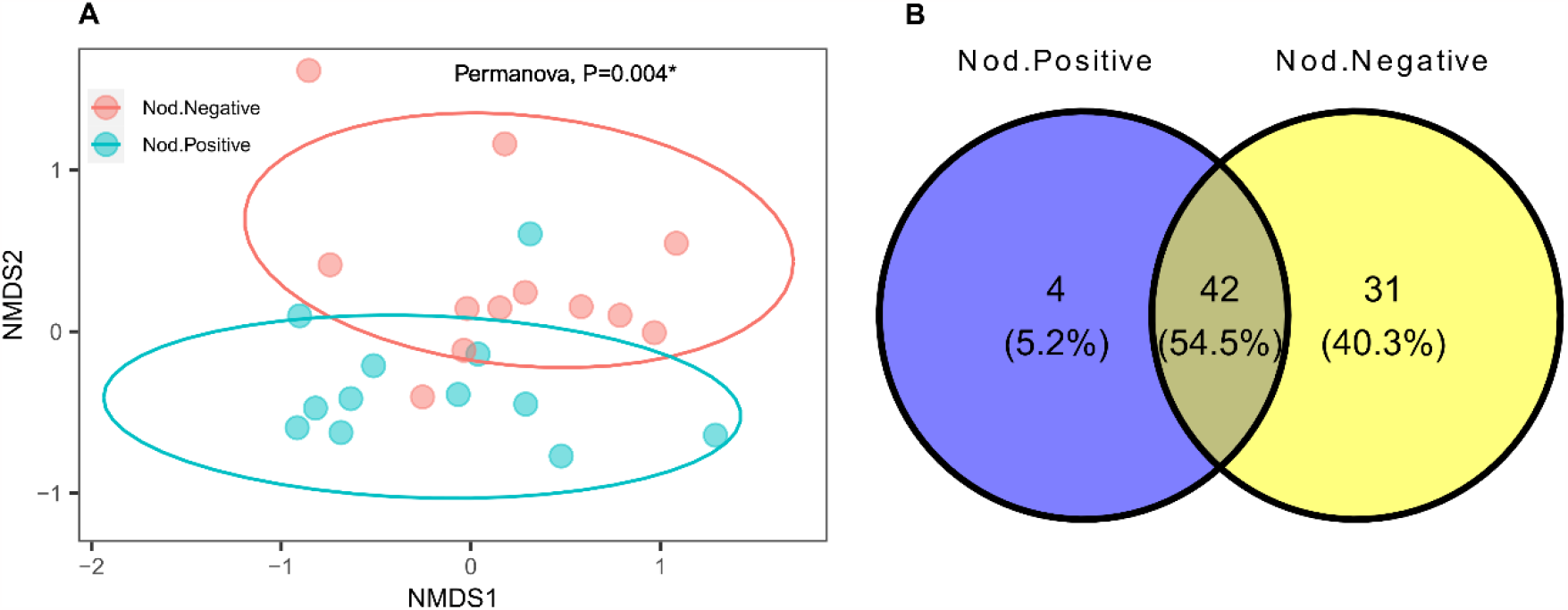
Panel A: NDMS analysis of the foliar fungal communities according to nodulation capacity of eight Fabaceae tree species. Panel B: Venn diagram showing the number of orders of endophytic fungi shared among all the tree species evaluated, and those found exclusively on positive nodulation plants or those plants that do not form nodules.

## 4. Discussion

In this study, we explore the diversity of fungal communities in the leaves of eight tropical tree species of the Fabaceae family. Our work is novel in demonstrating that the establishment, or not, of symbiotic relationships with nitrogen-fixing bacteria in roots has a profound effect on the richness and diversity of the mycobiome of their leaves. We highlight, in particular, that the diversity of fungal communities was significantly higher in plants that do not form nodules on their roots.

In general, there are few studies analyzing the effect of plant-bacterial relationships in one tissue (e.g., root) on plant-fungal relationships in another tissue (e.g., leaf), so it is difficult to elucidate the specific reasons for this phenomenon. However, we theorize that the high levels of fungal species richness and Shannon diversity detected in leaves of negative nodulation plants could be related to the principle of competitive exclusion, in this case by the availability of nitrogen (Hardin, 1960).

In nutrient-rich environments, competition for resources can lead to rapid dominance of a few favored species, outcompeting others and reducing overall diversity (Younginger et al., 2023). This effect is potentially intensified in nitrogen-fixing leguminous plants where higher nitrogen levels may stimulate the growth of certain fungi and displace others(Castellanos et al., 2018; Dovrat et al., 2020). Conversely, environmental constraints can drive the evolution of strategies for distinct resource use or the occupation of specific niches in the habitat, facilitating coexistence by reducing direct competition (Miller and Rudgers, 2014). This evolutionary process can generate distinct niches and increase diversity with each species occupying an ecological niche and adopting diverse strategies for resource acquisition and usage (Gerz et al., 2018; Pimm, 1983).

Nitrogen-fixing legumes may benefit from a more stable and direct source of nitrogen via root nodules. This potentially reduces their reliance on diverse endophytic fungi for nutrient acquisition and processing (Adeleke and Babalola, 2021; Behie et al., 2015). In contrast, plant species lacking the ability to fix nitrogen may depend more heavily on nutrient acquisition, potentially favoring a higher diversity of fungi with complementary uptake and processing capabilities.

In environments limited in essential nutrients like nitrogen, mutualistic relationships between fungal species can increase diversity. This occurs because different fungal species depend on each other for essential nutrients and resources, promoting a symbiotic network that benefits all involved (Dal Bello et al., 2021; Jansa et al., 2013; Jones et al., 2012). These mutualisms ultimately enhance the fitness of each species, potentially protecting them from competitive exclusion and favoring the coexistence of potential competitors (Johnson, 2021; Sprent, 2009).

In addition, plants of the Fabaceae family can recruit different types of soil microorganisms according to their ability to form nodules or not (Chang et al., 2017; Lagunas et al., 2023); thus, not only particular groups of nitrogen-fixing bacteria are associated with certain legume species, but the presence of a nitrogen-fixing symbiosis is directly involved in the selection of specific microbial taxa and changes in the host community (Lagunas et al., 2023; Mahmud et al., 2020; Trivedi et al., 2020; Zgadzaj et al., 2016).

Another reason that could explain the observed differences would be the presence of antifungal compounds in leaves as a result of symbiosis with nitrogen-fixing bacteria (Chaparro et al., 2014). In this sense, root exudates can vary their type and composition due to the establishment of symbiotic interactions with rhizobacteria, modulating at the same time the expression of genes involved in the synthesis of antifungal compounds (Mavrodi et al., 2021; Mohammed et al., 2020; Nóbrega et al., 2005; Xia et al., 2021).

The composition of the mycobiome is not solely driven by external factors but is also strongly controlled by the plant itself. Several chemical compounds, secondary metabolites, mucilage, and nutrients present in the root exudate can act as a selective force, attracting and favoring specific microorganisms. These selected microorganisms then migrate from the root to different plant organs, with the leaves being the main destination (Chaparro et al., 2014; Ferguson et al., 2019; Lagunas et al., 2023; Sánchez-Cañizares et al., 2017).

The endophytic fungal community of the analyzed plant species, regardless of their nodulation capacity, was dominated by the phylum Ascomycota. This observation aligns with previous research on other tropical plant species, where Ascomycota has consistently been found to be the most abundant phylum. The dominance of Ascomycota is likely due to their diverse metabolic capabilities, enabling them to thrive in various ecological niches and establish beneficial relationships with their host plants (Arnold and Lutzoni, 2007; Douanla-Meli and Langer, 2012; Sadeghi et al., 2019; Zhou et al., 2017).

While many endophytic ascomycetes exhibit generalist tendencies, they can also demonstrate remarkable tissue specificity within the same host plant. This means that specific fungal species may be preferentially found in certain plant organs, such as roots or leaves, even though they possess the ability to colonize other parts of the plant. This tissue-specific distribution suggests that endophytic ascomycetes may have evolved adaptations to optimize their interactions with different plant tissues and fulfill distinct ecological functions within the host (Rojas-Jimenez et al., 2016; Rojas-Jimenez and Tamayo-Castillo, 2021; Silva et al., 2023).

The prevalence of fungal orders Pleosporales and Capnodiales, represented by genera such as *Preussia, Corynespora*, and *Cladosporium*, underscores the presence of generalist endophytes across the analyzed plant species. This observation holds regardless of the host’s nodulation capacity or subfamily within the Fabaceae family. This widespread distribution likely reflects the remarkable functional versatility and diverse lifestyles of these fungal groups, allowing them to thrive in and benefit various plant species and tissues (dos Reis et al., 2022; Mani et al., 2015; Pietro-Souza et al., 2017).

This study reveals that interactions of Fabaceae plants with nitrogen-fixing bacteria in roots have a substantial impact on the richness and diversity of the mycobiome present in leaves. This finding suggests that dynamics at the root level can significantly influence the composition of fungal communities in other plant tissues(Lagunas et al., 2023; Mahmud et al., 2020). The lack of research addressing the influence of specific fungal orders in determining diversity based on nodulation capacity highlights the need for further exploration of these orders and their relationship to the establishment of plant-microorganism symbiosis.

To better understand the tripartite relationships between leguminous plants, root nitrogen-fixing bacteria, and endophytic fungi, future work should include complementary studies such as chemical analysis of elements in plant tissues, evaluation of the presence of mycorrhizae, and analysis of the chemical composition of root exudates as well as other metabolites in leaves. These studies could be done both in greenhouses and under natural conditions considering a larger number of samples.

Our work contributes to a better understanding of complex plant-microorganism interactions, particularly between tropical Fabaceae tree species and their endophytic fungi. By highlighting the influence of nodulation capacity on the configuration of leaf mycobiome, we provide insight into how root-level interactions can have cascading effects on other parts of the plant. These findings could also have practical applications in plant health improvement, agriculture, and natural ecosystem management.

## Data availability

Sequence data were deposited at the NCBI Sequence Read Archive under accession number PRJNA1048874 (https://www.ncbi.nlm.nih.gov/sra/PRJNA1048874).

## Acknowledgments

Funding was provided by University of Costa Rica, project C0-524. We thank Eduardo Chacón for reviewing the manuscript.

## References

Adams, M.A., Turnbull, T.L., Sprent, J.I., Buchmann, N., 2016. Legumes are different: Leaf nitrogen, photosynthesis, and water use efficiency. Proc. Natl. Acad. Sci. 113, 4098–4103. 10.1073/pnas.1523936113

Adeleke, B.S., Babalola, O.O., 2021. Biotechnological overview of agriculturally important endophytic fungi. Hortic. Environ. Biotechnol. 62, 507–520. 10.1007/s13580-021-00334-1

Alam, B., Lǐ, J., Gě, Q., Khan, M.A., Gōng, J., Mehmood, S., Yuán, Y., Gǒng, W., 2021. Endophytic Fungi: From Symbiosis to Secondary Metabolite Communications or Vice Versa?. Front. Plant Sci..

Arnold, A.E., Lutzoni, F., 2007. Diversity and host range of foliar fungal endophytes: are tropical leaves biodiversity hotspots? Ecology 88, 541–549.

Arnold, A.E., Mejía, L.C., Kyllo, D., Rojas, E.I., Maynard, Z., Robbins, N., Herre, E.A., 2003. Fungal endophytes limit pathogen damage in a tropical tree. Proc. Natl. Acad. Sci. 100, 15649–15654. 10.1073/pnas.2533483100

Azani, N., Babineau, M., Bailey, C.D., Banks, H., Barbosa, A.R., Pinto, R.B., Boatwright, J.S., Borges, L.M., Brown, G.K., Bruneau, A., Candido, E., Cardoso, D., Chung, K.-F., Clark, R.P., Conceição, A. de S., Crisp, M., Cubas, P., Delgado-Salinas, A., Dexter, K.G., Doyle, J.J., Duminil, J., Egan, A.N., de la Estrella, M., Falcão, M.J., Filatov, D.A., Fortuna-Perez, A.P., Fortunato, R.H., Gagnon, E., Gasson, P., Rando, J.G., de Azevedo Tozzi, A.M.G., Gunn, B., Harris, D., Haston, E., Hawkins, J.A., Herendeen, P.S., Hughes, C.E., Iganci, J.R. V, Javadi, F., Kanu, S.A., Kazempour-Osaloo, S., Kite, G.C., Klitgaard, B.B., Kochanovski, F.J., Koenen, E.J.M., Kovar, L., Lavin, M., le Roux, M., Lewis, G.P., de Lima, H.C., López-Roberts, M.C., Mackinder, B., Maia, V.H., Malécot, V., Mansano, V.F., Marazzi, B., Mattapha, S., Miller, J.T., Mitsuyuki, C., Moura, T., Murphy, D.J., Nageswara-Rao, M., Nevado, B., Neves, D., Ojeda, D.I., Pennington, R.T., Prado, D.E., Prenner, G., de Queiroz, L.P., Ramos, G., Filardi, F.L.R., Ribeiro, P.G., de Lourdes Rico-Arce, M., Sanderson, M.J., Santos-Silva, J., São-Mateus, W.M.B., Silva, M.J.S., Simon, M.F., Sinou, C., Snak, C., de Souza, É.R., Sprent, J., Steele, K.P., Steier, J.E., Steeves, R., Stirton, C.H., Tagane, S., Torke, B.M., Toyama, H., da Cruz, D.T., Vatanparast, M., Wieringa, J.J., Wink, M., Wojciechowski, M.F., Yahara, T., Yi, T., Zimmerman, E., 2017. A new subfamily classification of the Leguminosae based on a taxonomically comprehensive phylogeny: The Legume Phylogeny Working Group (LPWG). Taxon 66, 44–77. 10.12705/661.3

Behie, S.W., Jones, S.J., Bidochka, M.J., 2015. Plant tissue localization of the endophytic insect pathogenic fungi Metarhizium and Beauveria. Fungal Ecol. 13, 112–119. 10.1016/J.FUNECO.2014.08.001

Callahan, B.J., McMurdie, P.J., Rosen, M.J., Han, A.W., Johnson, A.J.A., Holmes, S.P., 2016. DADA2: High-resolution sample inference from Illumina amplicon data. Nat. Methods 13, 581–583. 10.1038/nmeth.3869

Castellanos, A.E., Llano-Sotelo, J.M., Machado-Encinas, L.I., López-Piña, J.E., Romo-Leon, J.R., Sardans, J., Peñuelas, J., 2018. Foliar C, N, and P stoichiometry characterize successful plant ecological strategies in the Sonoran Desert. Plant Ecol. 219, 775–788. 10.1007/s11258-018-0833-3

Chang, C., Nasir, F., Ma, L., Tian, C., 2017. Molecular Communication and Nutrient Transfer of Arbuscular Mycorrhizal Fungi, Symbiotic Nitrogen-Fixing Bacteria, and Host Plant in Tripartite Symbiosis, in: Legume Nitrogen Fixation in Soils with Low Phosphorus Availability. Springer International Publishing, Cham, pp. 169–183. 10.1007/978-3-319-55729-8_9

Chaparro, J.M., Badri, D. V, Vivanco, J.M., 2014. Rhizosphere microbiome assemblage is affected by plant development. ISME J. 8, 790–803. 10.1038/ismej.2013.196

Clay, K., Schardl, C., 2002. Evolutionary origins and ecological consequences of endophyte symbiosis with grasses. Am. Nat. 160, 99–127.

Dal Bello, M., Lee, H., Goyal, A., Gore, J., 2021. Resource–diversity relationships in bacterial communities reflect the network structure of microbial metabolism. Nat. Ecol. Evol. 5, 1424–1434. 10.1038/s41559-021-01535-8

Danko, D., Bezdan, D., Afshin, E.E., Ahsanuddin, S., Bhattacharya, C., Butler, D.J., Chng, K.R., Donnellan, D., Hecht, J., Jackson, K., Kuchin, K., Karasikov, M., Lyons, A., Mak, L., Meleshko, D., Mustafa, H., Mutai, B., Neches, R.Y., Ng, A., Nikolayeva, O., Nikolayeva, T., Png, E., Ryon, K.A., Sanchez, J.L., Shaaban, H., Sierra, M.A., Thomas, D., Young, B., Abudayyeh, O.O., Alicea, J., Bhattacharyya, M., Blekhman, R., Castro-Nallar, E., Cañas, A.M., Chatziefthimiou, A.D., Crawford, R.W., De Filippis, F., Deng, Y., Desnues, C., Dias-Neto, E., Dybwad, M., Elhaik, E., Ercolini, D., Frolova, A., Gankin, D., Gootenberg, J.S., Graf, A.B., Green, D.C., Hajirasouliha, I., Hastings, J.J.A., Hernandez, M., Iraola, G., Jang, S., Kahles, A., Kelly, F.J., Knights, K., Kyrpides, N.C., Łabaj, P.P., Lee, P.K.H., Leung, M.H.Y., Ljungdahl, P.O., Mason-Buck, G., McGrath, K., Meydan, C., Mongodin, E.F., Moraes, M.O., Nagarajan, N., Nieto-Caballero, M., Noushmehr, H., Oliveira, M., Ossowski, S., Osuolale, O.O., Özcan, O., Paez-Espino, D., Rascovan, N., Richard, H., Rätsch, G., Schriml, L.M., Semmler, T., Sezerman, O.U., Shi, L., Shi, T., Siam, R., Song, L.H., Suzuki, H., Court, D.S., Tighe, S.W., Tong, X., Udekwu, K.I., Ugalde, J.A., Valentine, B., Vassilev, D.I., Vayndorf, E.M., Velavan, T.P., Wu, J., Zambrano, M.M., Zhu, J., Zhu, S., Mason, C.E., Abdullah, N., Abraao, M., Adel, A., Afaq, M., Al-Quaddoomi, F.S., Alam, I., Albuquerque, G.E., Alexiev, A., Ali, K., Alvarado-Arnez, L.E., Aly, S., Amachee, J., Amorim, M.G., Ampadu, M., Amran, M.A.-F., An, N., Andrew, W., Andrianjakarivony, H., Angelov, M., Antelo, V., Aquino, C., Aranguren, Á., Araujo, L.F., Vasquez Arevalo, H.F., Arevalo, J., Arnan, C., Alvarado Arnez, L.E., Arredondo, F., Arthur, M., Asenjo, F., Aung, T.S., Auvinet, J., Aventin, N., Ayaz, S., Baburyan, S., Bakere, A.-M., Bakhl, K., Bartelli, T.F., Batdelger, E., Baudon, F., Becher, K., Bello, C., Benchouaia, M., Benisty, H., Benoiston, A.-S., Benson, J., Benítez, D., Bernardes, J., Bertrand, D., Beurmann, S., Bitard-Feildel, T., Bittner, L., Black, C., Blanc, G., Blyther, B., Bode, T., Boeri, J., Boldgiv, B., Bolzli, K., Bordigoni, A., Borrelli, C., Bouchard, S., Bouly, J.-P., Boyd, A., Branco, G.P., Breschi, A., Brindefalk, B., Brion, C., Briones, A., Buczansla, P., Burke, C.M., Burrell, A., Butova, A., Buttar, I., Bynoe, J., Bönigk, S., Bøifot, K.O., Caballero, H., Cai, X.W., Calderon, D., Cantillo, A., Carbajo, M., Carbone, A., Cardenas, A., Carrillo, K., Casalot, L., Castro, S., Castro, Ana V., Castro, Astred, Castro, A.V.B., Cawthorne, S., Cedillo, J., Chaker, S., Chalangal, J., Chan, A., Chasapi, A.I., Chatziefthimiou, S., Chaudhuri, S.R., Chavan, A.K., Chavez, F., Chem, G., Chen, X., Chen, M., Chen, J.-W., Chernomoretz, A., Chettouh, A., Cheung, D., Chicas, D., Chiu, S., Choudhry, H., Chrispin, C., Ciaramella, K., Cifuentes, E., Cohen, J., Coil, D.A., Collin, S., Conger, C., Conte, R., Corsi, F., Cossio, C.N., Costa, A.F., Cuebas, D., D’Alessandro, B., Dahlhausen, K.E., Darling, A.E., Das, P., Davenport, L.B., David, L., Davidson, N.R., Dayama, G., Delmas, S., Deng, C.K., Dequeker, C., Desert, A., Devi, M., Dezem, F.S., Dias, C.N., Donahoe, T.R., Dorado, S., Dorsey, L., Dotsenko, V., Du, S., Dutan, A., Eady, N., Eisen, J.A., Elaskandrany, M., Epping, L., Escalera-Antezana, J.P., Ettinger, C.L., Faiz, I., Fan, L., Farhat, N., Faure, E., Fauzi, F., Feigin, C., Felice, S., Ferreira, L.P., Figueroa, G., Fleiss, A., Flores, D., Velasco Flores, J.L., Fonseca, M.A.S., Foox, J., Forero, J.C., Francis, A., French, K., Fresia, P., Friedman, J., Fuentes, J.J., Galipon, J., Garcia, M., Garcia, L., García, C., Geiger, A., Gerner, S.M., Ghose, S.L., Giang, D.P., Giménez, M., Giovannelli, D., Githae, D., Gkotzis, S., Godoy, L., Goldman, S., Gonnet, G.H., Gonzalez, J., Gonzalez, A., Gonzalez-Poblete, C., Gray, A., Gregory, T., Greselle, C., Guasco, S., Guerra, J., Gurianova, N., Haehr, W., Halary, S., Hartkopf, F., Hastings, J.J.A., Hawkins-Zafarnia, A., Hazrin-Chong, N.H., Helfrich, E., Hell, E., Henry, T., Hernandez, S., Hernandez, P.L., Hess-Homeier, D., Hittle, L.E., Hoan, N.X., Holik, A., Homma, C., Hoxie, I., Huber, M., Humphries, E., Hyland, S., Hässig, A., Häusler, R., Hüsser, N., Petit, R.A., Iderzorig, B., Igarashi, M., Iqbal, S.B., Ishikawa, S., Ishizuka, S., Islam, S., Islam, R., Ito, K., Ito, S., Ito, T., Ivankovic, T., Iwashiro, T., Jackson, S., Jacobs, J., James, M., Jaubert, M., Jerier, M.-L., Jiminez, E., Jinfessa, A., De Jong, Y., Joo, H.W., Jospin, G., Kajita, T., Ahmad Kassim, A.S., Kato, N., Kaur, A., Kaur, I., de Souza Gomes Kehdy, F., Khadka, V.S., Khan, S., Khavari, M., Ki, M., Kim, G., Kim, H.J., Kim, S., King, R.J., Knights, K., KoLoMonaco, G., Koag, E., Kobko-Litskevitch, N., Korshevniuk, M., Kozhar, M., Krebs, J., Kubota, N., Kuklin, A., Kumar, S.S., Kwong, R., Kwong, L., Lafontaine, I., Lago, J., Lai, T.Y., Laine, E., Laiola, M., Lakhneko, O., Lamba, I., de Lamotte, G., Lannes, R., De Lazzari, E., Leahy, M., Lee, H., Lee, Y., Lee, L., Lemaire, V., Leong, E., Leung, M.H.Y., Lewandowska, D., Li, C., Liang, W., Lin, M., Lisboa, P., Litskevitch, A., Liu, E.M., Liu, T., Livia, M.A., Lo, Y.H., Losim, S., Loubens, M., Lu, J., Lykhenko, O., Lysakova, S., Mahmoud, S., Majid, S.A., Makogon, N., Maldonado, D., Mallari, K., Malta, T.M., Mamun, M., Manoir, D., Marchandon, G., Marciniak, N., Marinovic, S., Marques, B., Mathews, N., Matsuzaki, Y., Matthys, V., May, M., McComb, E., Meagher, A., Melamed, A., Menary, W., Mendez, K.N., Mendez, A., Mendy, I.M., Meng, I., Menon, A., Menor, M., Meoded, R., Merino, N., Meydan, C., Miah, K., Mignotte, M., Miketic, T., Miranda, W., Mitsios, A., Miura, R., Miyake, K., Moccia, M.D., Mohan, N., Mohsin, M., Moitra, K., Moldes, M., Molina, L., Molinet, J., Molomjamts, O.-E., Moniruzzaman, E., Moon, S., de Oliveira Moraes, I., Moreno, M., Mosella, M.S., Moser, J.W., Mozsary, C., Muehlbauer, A.L., Muner, O., Munia, M., Munim, N., Muscat, M., Mustac, T., Muñoz, C., Nadalin, F., Naeem, A., Nagy-Szakal, D., Nakagawa, M., Narce, A., Nasu, M., Navarrete, I.G., Naveed, H., Nazario, B., Nedunuri, N.R., Neff, T., Nesimi, A., Ng, W.C., Ng, S., Nguyen, G., Ngwa, E., Nicolas, A., Nicolas, P., Nika, A., Noorzi, H., Nosrati, A., Noushmehr, H., Nunes, D.N., O’Brien, K., O’Hara, N.B., Oken, G., Olawoyin, R.A., Oliete, J.Q., Olmeda, K., Oluwadare, T., Oluwadare, I.A., Ordioni, N., Orpilla, J., Orrego, J., Ortega, M., Osma, P., Osuolale, I.O., Osuolale, O.M., Ota, M., Oteri, F., Oto, Y., Ounit, R., Ouzounis, C.A., Pakrashi, S., Paras, R., Pardo-Este, C., Park, Y.-J., Pastuszek, P., Patel, S., Pathmanathan, J., Patrignani, A., Perez, M., Peros, A., Persaud, S., Peters, A., Phillips, A., Pineda, L., Pizzi, M.P., Plaku, Alma, Plaku, Alketa, Pompa-Hogan, B., Portilla, M.G., Posada, L., Priestman, M., Prithiviraj, B., Priya, S., Pugdeethosal, P., Pugh, C.E., Pulatov, B., Pupiec, A., Pyrshev, K., Qing, T., Rahiel, S., Rahmatulloev, S., Rajendran, K., Ramcharan, A., Ramirez-Rojas, A., Rana, S., Ratnanandan, P., Read, T.D., Rehrauer, H., Richer, R., Rivera, A., Rivera, M., Robertiello, A., Robinson, C., Rodríguez, P., Rojas, N.A., Roldán, P., Rosario, A., Roth, S., Ruiz, M., Boja Ruiz, S.E., Russell, K., Rybak, M., Sabedot, T.S., Sabina, M., Saito, I., Saito, Y., Malca Salas, G.A., Salazar, C., San, K.M., Sanchez, J., Sanchir, K., Sankar, R., de Souza Santos, P.T., Saravi, Z., Sasaki, K., Sato, Y., Sato, M., Sato, S., Sato, R., Sato, K., Sayara, N., Schaaf, S., Schacher, O., Schinke, A.-L.M., Schlapbach, R., Schori, C., Schriml, J.R., Segato, F., Sepulveda, F., Serpa, M.S., De Sessions, P.F., Severyn, J.C., Shaaban, H., Shakil, M., Shalaby, S., Shari, A., Shim, H., Shirahata, H., Shiwa, Y., Siam, R., Da Silva, O., Silva, J.M., Simon, G., Singh, S.K., Sluzek, K., Smith, R., So, E., Andreu Somavilla, N., Sonohara, Y., Rufino de Sousa, N., Souza, C., Sperry, J., Sprinsky, N., Stark, S.G., La Storia, A., Suganuma, K., Suliman, H., Sullivan, J., Supie, A.A.M., Suzuki, C., Takagi, S., Takahara, F., Takahashi, N., Takahashi, K., Takeda, T., Takenaka, I.K., Tanaka, S., Tang, A., Man Tang, Y., Tarcitano, E., Tassinari, A., Taye, M., Terrero, A., Thambiraja, E., Thiébaut, A., Thomas, S., Thomas, A.M., Togashi, Y., Togashi, T., Tomaselli, A., Tomita, M., Tomita, I., Tong, X., Toth, O., Toussaint, N.C., Tran, J.M., Truong, C., Tsonev, S.I., Tsuda, K., Tsurumaki, T., Tuz, M., Tymoshenko, Y., Urgiles, C., Usui, M., Vacant, S., Valentine, B., Vann, L.E., Velter, F., Ventorino, V., Vera-Wolf, P., Vicedomini, R., Suarez-Villamil, M.A., Vincent, S., Vivancos-Koopman, R., Wan, A., Wang, C., Warashina, T., Watanabe, A., Weekes, S., Werner, J., Westfall, D., Wieler, L.H., Williams, M., Wolf, S.A., Wong, B., Wong, Y.L., Wong, T., Wright, R., Wunderlin, T., Yamanaka, R., Yang, J., Yano, H., Yeh, G.C., Yemets, O., Yeskova, T., Yoshikawa, S., Zafar, L., Zhang, Y., Zhang, S., Zhang, A., Zheng, Y., Zubenko, S., 2021. A global metagenomic map of urban microbiomes and antimicrobial resistance. Cell. 10.1016/j.cell.2021.05.002

de Bedout-Mora, M., Solis-Ramos, L., Valverde-Barrantes, O., Rojas-Jiménez, K., 2022. Capacidad de nodulación en especies forestales leguminosas (Fabaceae) según su filogenia y características morfológicas. Rev. For. Mesoam. Kurú 19. 10.18845/rfmk.v19i45.6315

de Faria, S.M., Ringelberg, J.J., Gross, E., Koenen, E.J.M., Cardoso, D., Ametsitsi, G.K.D., Akomatey, J., Maluk, M., Tak, N., Gehlot, H.S., Wright, K.M., Teaumroong, N., Songwattana, P., de Lima, H.C., Prin, Y., Zartman, C.E., Sprent, J.I., Ardley, J., Hughes, C.E., James, E.K., 2022. The innovation of the symbiosome has enhanced the evolutionary stability of nitrogen fixation in legumes. New Phytol. 235, 2365–2377. 10.1111/nph.18321

dos Reis, J.B.A., do Vale, H.M.M., Lorenzi, A.S., 2022. Insights into taxonomic diversity and bioprospecting potential of Cerrado endophytic fungi: a review exploring an unique Brazilian biome and methodological limitations. World J. Microbiol. Biotechnol. 38, 202. 10.1007/s11274-022-03386-2

Douanla-Meli, C., Langer, E., 2012. Diversity and molecular phylogeny of fungal endophytes associated with Diospyros crassiflora. Mycology 3, 175–187. 10.1080/21501203.2012.705348

Dovrat, G., Bakhshian, H., Masci, T., Sheffer, E., 2020. The nitrogen economic spectrum of legume stoichiometry and fixation strategy. New Phytol. 227, 365–375. 10.1111/nph.16543

Ferguson, B.J., Indrasumunar, A., Hayashi, S., Lin, M.-H., Lin, Y., Reid, D.E., Gresshoff, P.M., 2010. Molecular Analysis of Legume Nodule Development and Autoregulation. J. Integr. Plant Biol. 52, 61–76. 10.1111/j.1744-7909.2010.00899.x

Ferguson, B.J., Mens, C., Hastwell, A.H., Zhang, M., Su, H., Jones, C.H., Chu, X., Gresshoff, P.M., 2019. Legume nodulation: The host controls the party. Plant. Cell Environ. 42, 41–51. 10.1111/pce.13348

Franklin, J.B., Hockey, K., Maherali, H., 2020. Population‐level variation in host plant response to multiple microbial mutualists. Am. J. Bot. 107, 1389–1400. 10.1002/ajb2.1543

Gerz, M., Guillermo Bueno, C., Ozinga, W.A., Zobel, M., Moora, M., 2018. Niche differentiation and expansion of plant species are associated with mycorrhizal symbiosis. J. Ecol. 106, 254–264. 10.1111/1365-2745.12873

Hardin, G., 1960. The Competitive Exclusion Principle. Science (80-.). 131, 1292–1297. 10.1126/science.131.3409.1292

Heilmann-Clausen, J., Maruyama, P.K., Bruun, H.H., Dimitrov, D., Læssøe, T., Frøslev, T.G., Dalsgaard, B., 2016. Citizen science data reveal ecological, historical and evolutionary factors shaping interactions between woody hosts and wood-inhabiting fungi. New Phytol. 212, 1072–1082. 10.1111/nph.14194

James, E.K., 2022. The seeds of nodulation. J. Plant Physiol. 278, 153812. 10.1016/j.jplph.2022.153812

Jansa, J., Bukovská, P., Gryndler, M., 2013. Mycorrhizal hyphae as ecological niche for highly specialized hypersymbionts – or just soil free-riders? Front. Plant Sci. 4. 10.3389/fpls.2013.00134

Johnson, C.A., 2021. How mutualisms influence the coexistence of competing species. Ecology 102. 10.1002/ecy.3346

Jones, E.I., Bronstein, J.L., Ferrière, R., 2012. The fundamental role of competition in the ecology and evolution of mutualisms. Ann. N. Y. Acad. Sci. 1256, 66–88. 10.1111/j.1749-6632.2012.06552.x

Lagunas, B., Richards, L., Sergaki, C., Burgess, J., Pardal, A.J., Hussain, R.M.F., Richmond, B.L., Baxter, L., Roy, P., Pakidi, A., Stovold, G., Vázquez, S., Ott, S., Schäfer, P., Gifford, M.L., 2023. Rhizobial nitrogen fixation efficiency shapes endosphere bacterial communities and Medicago truncatula host growth. Microbiome 11, 146. 10.1186/s40168-023-01592-0

Lewis, G.P., 2005. Legumes of the world, TA - TT -. Royal Botanic Gardens, Kew Richmond, UK, Richmond, UK SE - xiv, 577 pages : illustrations (some color), maps (some color) ; 30 cm.

Mahmud, K., Makaju, S., Ibrahim, R., Missaoui, A., 2020. Current Progress in Nitrogen Fixing Plants and Microbiome Research. Plants 9, 97. 10.3390/plants9010097

Mani, V.M., Gnana Soundari, A.P., Karthiyaini, D., Preethi, K., 2015. Bioprospecting Endophytic Fungi and Their Metabolites from Medicinal Tree Aegle marmelos in Western Ghats, India. Mycobiology 43, 303–310. 10.5941/MYCO.2015.43.3.303

Mathesius, U., 2022. Are legumes different? Origins and consequences of evolving nitrogen fixing symbioses. J. Plant Physiol. 276, 153765. 10.1016/j.jplph.2022.153765

Mavrodi, O. V., McWilliams, J.R., Peter, J.O., Berim, A., Hassan, K.A., Elbourne, L.D.H., LeTourneau, M.K., Gang, D.R., Paulsen, I.T., Weller, D.M., Thomashow, L.S., Flynt, A.S., Mavrodi, D. V., 2021. Root Exudates Alter the Expression of Diverse Metabolic, Transport, Regulatory, and Stress Response Genes in Rhizosphere Pseudomonas. Front. Microbiol. 12. 10.3389/fmicb.2021.651282

Miller, T.E.X., Rudgers, J.A., 2014. Niche Differentiation in the Dynamics of Host-Symbiont Interactions: Symbiont Prevalence as a Coexistence Problem. Am. Nat. 183, 506–518. 10.1086/675394

Mohammed, H., Jaiswal, S.K., Mohammed, M., Mbah, G.C., Dakora, F.D., 2020. Insights into nitrogen fixing traits and population structure analyses in cowpea (Vigna unguiculata L. Walp) accessions grown in Ghana. Physiol. Mol. Biol. Plants 26, 1263–1280. 10.1007/s12298-020-00811-4

Molina, R., 1992. Specificity Phenomena in Mycorrhizal Symbioses: Community-Ecological Consequences and Practical Implications Randy Molina, Hugues Massicotte, and James M. Trappe. Mycorrhizal functioning: an integrative plant-fungal process. Mycorrhizal Funct. an Integr. plant-fungal Process 357.

Nóbrega, F.M., Santos, I.S., Cunha, M.D., Carvalho, A.O., Gomes, V.M., 2005. Antimicrobial proteins from cowpea root exudates: inhibitory activity against Fusarium oxysporum and purification of a chitinase-like protein. Plant Soil 272, 223–232. 10.1007/s11104-004-4954-1

Oksanen, J., Blanchet, F.G., Friendly, M., Kindt, R., Legendre, P., McGlinn, D., Minchin, P.R., O’Hara, R.B., Simpson, G.L., Solymos, P., Stevens, M.H.H., Szoecs, E., Wagner, H., 2022. vegan: Community Ecology Package.

Parker, M.A., 2008. Symbiotic Relationships of Legumes and Nodule Bacteria on Barro Colorado Island, Panama: A Review. Microb. Ecol. 55, 662–672. 10.1007/s00248-007-9309-z

Pietro-Souza, W., Mello, I.S., Vendruscullo, S.J., Silva, G.F. da, Cunha, C.N. da, White, J.F., Soares, M.A., 2017. Endophytic fungal communities of Polygonum acuminatum and Aeschynomene fluminensis are influenced by soil mercury contamination. PLoS One 12, e0182017. 10.1371/journal.pone.0182017

Pimm, S.L., 1983. Tilman, D. 1982. Resource competition and community structure. Monogr. Pop. Biol. 17. Princeton University Press, Princeton, N.J. 296 p. $27.50. Limnol. Oceanogr. 28, 1043–1045. 10.4319/lo.1983.28.5.1043

Põlme, S., Abarenkov, K., Henrik Nilsson, R., Lindahl, B.D., Clemmensen, K.E., Kauserud, H., Nguyen, N., Kjøller, R., Bates, S.T., Baldrian, P., Frøslev, T.G., Adojaan, K., Vizzini, A., Suija, A., Pfister, D., Baral, H.-O., Järv, H., Madrid, H., Nordén, J., Liu, J.-K., Pawlowska, J., Põldmaa, K., Pärtel, K., Runnel, K., Hansen, K., Larsson, K.-H., Hyde, K.D., Sandoval-Denis, M., Smith, M.E., Toome-Heller, M., Wijayawardene, N.N., Menolli, N., Reynolds, N.K., Drenkhan, R., Maharachchikumbura, S.S.N., Gibertoni, T.B., Læssøe, T., Davis, W., Tokarev, Y., Corrales, A., Soares, A.M., Agan, A., Machado, A.R., Argüelles-Moyao, A., Detheridge, A., de Meiras-Ottoni, A., Verbeken, A., Dutta, A.K., Cui, B.-K., Pradeep, C.K., Marín, C., Stanton, D., Gohar, D., Wanasinghe, D.N., Otsing, E., Aslani, F., Griffith, G.W., Lumbsch, T.H., Grossart, H.-P., Masigol, H., Timling, I., Hiiesalu, I., Oja, J., Kupagme, J.Y., Geml, J., Alvarez-Manjarrez, J., Ilves, K., Loit, K., Adamson, K., Nara, K., Küngas, K., Rojas-Jimenez, K., Bitenieks, K., Irinyi, L., Nagy, L.L., Soonvald, L., Zhou, L.-W., Wagner, L., Aime, M.C., Öpik, M., Mujica, M.I., Metsoja, M., Ryberg, M., Vasar, M., Murata, M., Nelsen, M.P., Cleary, M., Samarakoon, M.C., Doilom, M., Bahram, M., Hagh-Doust, N., Dulya, O., Johnston, P., Kohout, P., Chen, Q., Tian, Q., Nandi, R., Amiri, R., Perera, R.H., dos Santos Chikowski, R., Mendes-Alvarenga, R.L., Garibay-Orijel, R., Gielen, R., Phookamsak, R., Jayawardena, R.S., Rahimlou, S., Karunarathna, S.C., Tibpromma, S., Brown, S.P., Sepp, S.-K., Mundra, S., Luo, Z.-H., Bose, T., Vahter, T., Netherway, T., Yang, T., May, T., Varga, T., Li, W., Coimbra, V.R.M., de Oliveira, V.R.T., de Lima, V.X., Mikryukov, V.S., Lu, Y., Matsuda, Y., Miyamoto, Y., Kõljalg, U., Tedersoo, L., 2020. FungalTraits: a user-friendly traits database of fungi and fungus-like stramenopiles. Fungal Divers. 105, 1–16. 10.1007/s13225-020-00466-2

Põlme, S., Bahram, M., Jacquemyn, H., Kennedy, P., Kohout, P., Moora, M., Oja, J., Öpik, M., Pecoraro, L., Tedersoo, L., 2018a. Host preference and network properties in biotrophic plant–fungal associations. New Phytol. 217, 1230–1239. 10.1111/nph.14895

Põlme, S., Bahram, M., Jacquemyn, H., Kennedy, P., Kohout, P., Moora, M., Oja, J., Öpik, M., Pecoraro, L., Tedersoo, L., 2018b. Host preference and network properties in biotrophic plant–fungal associations. New Phytol. 217, 1230–1239. 10.1111/nph.14895

R Core Team, 2023. R: A Language and Environment for Statistical Computing.

Reis, J.B.A. dos, 2022. Micobiota endofítica de seis espécies lenhosas do Cerrado : efeitos da planta hospedeira e da adição de nutrientes ao solo. Universidade de Brasília.

Rodriguez, R.J., White Jr., J.F., Arnold, A.E., Redman, R.S., 2009. Fungal endophytes: diversity and functional roles. New Phytol. 182, 314–330. 10.1111/j.1469-8137.2009.02773.x

Rojas-Jimenez, K., Hernandez, M., Blanco, J., Vargas, L.D., Acosta-Vargas, L.G., Tamayo, G., 2016. Richness of cultivable endophytic fungi along an altitudinal gradient in wet forests of Costa Rica. Fungal Ecol. 20, 124–131. 10.1016/j.funeco.2015.12.006

Rojas-Jimenez, K., Tamayo-Castillo, G., 2021. Fungal Endophytes and Their Bioactive Compounds in Tropical Forests of Costa Rica, in: Rosa, L.H. (Ed.), Neotropical Endophytic Fungi. Springer International Publishing, Cham, pp. 109–130. 10.1007/978-3-030-53506-3_6

Sadeghi, F., Samsampour, D., Seyahooei, M.A., Bagheri, A., Soltani, J., 2019. Diversity and Spatiotemporal Distribution of Fungal Endophytes Associated with Citrus reticulata cv. Siyahoo. Curr. Microbiol. 76, 279–289. 10.1007/s00284-019-01632-9

Sánchez-Cañizares, C., Jorrín, B., Poole, P.S., Tkacz, A., 2017. Understanding the holobiont: the interdependence of plants and their microbiome. Curr. Opin. Microbiol. 38, 188–196. 10.1016/j.mib.2017.07.001

Schulz, B., Boyle, C., 2005. The endophytic continuum. Mycol. Res. 109, 661–686. 10.1017/S095375620500273X

Silva, P.S., Royo, V.A., Valerio, H.M., Fernandes, E.G., Queiroz, M. V., Fagundes, M., 2023. Filtrates from cultures of endophytic fungi isolated from leaves of Copaifera oblongifolia (Fabaceae) affect germination and seedling development differently. Brazilian J. Biol. 83. 10.1590/1519-6984.242070

Sprent, J.I., 2009. Legume Nodulation. Wiley. 10.1002/9781444316384

Sprent, J.I., 2008. Evolution and diversity of legume symbiosis, in: Nitrogen-Fixing Leguminous Symbioses. Springer, pp. 1–21.

Sprent, J.I., Ardley, J., James, E.K., 2017. Biogeography of nodulated legumes and their nitrogen‐ fixing symbionts. New Phytol. 215, 40–56. 10.1111/nph.14474

Trivedi, P., Leach, J.E., Tringe, S.G., Sa, T., Singh, B.K., 2020. Plant–microbiome interactions: from community assembly to plant health. Nat. Rev. Microbiol. 18, 607–621. 10.1038/s41579-020-0412-1

Xia, M., Valverde‐Barrantes, O.J., Suseela, V., Blackwood, C.B., Tharayil, N., 2021. Coordination between compound‐specific chemistry and morphology in plant roots aligns with ancestral mycorrhizal association in woody angiosperms. New Phytol. 232, 1259–1271. 10.1111/nph.17561

Younginger, B.S., Stewart, N.U., Balkan, M.A., Ballhorn, D.J., 2023. Stable coexistence or competitive exclusion? Fern endophytes demonstrate rapid turnover favouring a dominant fungus. Mol. Ecol. 32, 244–257. 10.1111/mec.16732

Zgadzaj, R., Garrido-Oter, R., Jensen, D.B., Koprivova, A., Schulze-Lefert, P., Radutoiu, S., 2016. Root nodule symbiosis in Lotus japonicus drives the establishment of distinctive rhizosphere, root, and nodule bacterial communities. Proc. Natl. Acad. Sci. 113. 10.1073/pnas.1616564113

Zhou, Y.-K., Shen, X.-Y., Hou, C.-L., 2017. Diversity and antimicrobial activity of culturable fungi from fishscale bamboo (Phyllostachys heteroclada) in China. World J. Microbiol. Biotechnol. 33, 104. 10.1007/s11274-017-2267-9

